# Cholinergic interneurons as a novel target of CRF in the striatum that is spared by repeated stress

**DOI:** 10.1101/448571

**Authors:** Julia C. Lemos, Jung Hoon Shin, Anna E. Ingebretson, Lauren K. Dobbs, Veronica A. Alvarez

**Author notes:** Julia C. Lemos is now at the University of Minnesota, Department of Neuroscience.

## Abstract

Acute stressors can stimulate appetitive and exploratory behaviors, not just produce negative affect that impair performance. Corticotropin releasing factor (CRF), which is released in the brain in response to stress, acts on different targets and circuits to mediate both the negative and positive effects of stress. In the nucleus accumbens, CRF facilitates appetitive behavior through mechanisms not fully understood. Here we report that cholinergic interneurons (CINs) are a novel target for CRF actions in the striatum. CRF enhances the spontaneous firing via activation of CRF-type 1 receptors expressed on CINs. This causes the activation muscarinic acetylcholine receptors type 5, which mediate CRF potentiation of dopamine transmission in the striatum. Repeated stress selectively dampens some CRF functions but spare effect on CINs and changes CRF-R1 expression in a cell-specific manner. These data highlight the existence of diverse CRF targets within the striatum, which vary in their resilience to stress.

## Introduction

CRF is a neuropeptide neuromodulator that is often released in response to salient environmental stimuli including acute stressors (Cook, 2004; Holly et al., 2016; Merali et al., 2004; Ohmura et al., 2009; Wang et al., 2005). CRF has canonically been defined as the initiation factor of the hypothalamic-pituitary-adrenal (HPA) axis response (Bale, 2006; Bale and Vale, 2004). It is densely expressed in the paraventricular nucleus of the hypothalamus (PVN) and triggers the release of ACTH from the anterior pituitary by activation of CRF-type 1 receptors (CRF-R1)(Bale and Vale, 2004; Nestler et al., 2002). Both CRF and its two receptors, CRF-R1 and CRF-type 2 (CRF-R2) are widely distributed throughout the brain, beyond the hypothalamic area (Henckens et al., 2016; Justice et al., 2008; Kuhne et al., 2012; Steckler and Holsboer, 1999; Van Pett et al., 2000). There is a rich literature investigating the actions of CRF in extrahypothalamic areas such as the amygdala, bed nucleus of the stria terminalis (BNST), locus coeruleus and dorsal raphe nucleus (DRN). Historically, little attention has been paid to the striatal complex owing to the relatively low expression levels of both CRF neuropeptide and receptors compared to the aforementioned areas (Henckens et al., 2016; Steckler and Holsboer, 1999; Van Pett et al., 2000). However, recent evidence suggests that CRF acting in the NAc can promote appetitive/approach behaviors including cue-triggered operant behaviors, novelty exploration and social affiliation behavior (Lemos et al., 2012; Lim et al., 2007; Pecina et al., 2006). This behavioral effect of CRF has been linked to the *in vitro* observation that CRF can potentiate DA transmission in brain slices, suggesting CRF actions at the level of the dopamine terminals within the NAc. Further *in vivo* microdialysis experiments also suggest that CRF potentiates dopamine transmission in the NAc independently of actions at the cell soma (Chen et al., 2012; Lemos et al., 2012). However, the mechanisms underlying this effect of CRF in the striatum remain unclear.

Microdialysis data has shown that CRF infusion into the NAc increases acetylcholine levels in the striatum, in addition to DA (Chen et al., 2012). Over the last few years, we have gained further understanding on how acetylcholine can both trigger and modulate dopamine release through activation of nicotinic and muscarinic acetylcholine receptors respectively. Work from Dani and colleagues had demonstrated that electrically stimulated dopamine transients evoked in an *ex vivo* slice preparation were reduced following antagonism of β2-containing nAChRs (Zhang et al., 2004; Zhou et al., 2002). It was subsequently demonstrated that optogenetic stimulation of CINs within the dorsal or ventral striatum evokes dopamine transients, and that this signal could be completely abolished by the nicotinic antagonists DhβE or mecamylamine (Cachope et al., 2012; Shin et al., 2017; Threlfell et al., 2012). The kinetics of these events along with electron microscopy data supports a working model where β2-containing nAChRs localized along dopamine axons and terminals can depolarize these compartments and evoke DA release in response to acetylcholine release (Wonnacott et al., 2000). Thus, synchronous firing of CINs can evoke DA release independently of the ventral tegmental area (VTA) or substantia nigra (SN) dopamine neuronal activity. Further, activation of muscarinic acetylcholine receptor can modulate DA transmission. Gi/o-coupled M2 and M4 receptors are expressed on striatal CINs and act as autoreceptors, suppressing both acetylcholine transmission and in turn, dopamine transmission (Shin et al., 2015; Threlfell et al., 2012; Yan and Surmeier, 1996). In contrast, Gq-coupled M5 receptors localized to dopamine processes potentiates dopamine transmission (Schmidt et al., 2010; Shin et al., 2015; Zhang et al., 2002). Based on these facts, we reasoned that CRF could enhance DA transmission by activating striatal CINs. And indeed, an independent study demonstrated that stress-evoked dopamine transmission in the NAc was reduced by selective ablation of CINs in NAc (Laplante et al., 2013). Thus, we sought to test this hypothesis directly by measuring the effects of exogenously applied CRF on the firing of CINs within the dorsal striatum (DS) and NAc. Moreover, we tested whether the CRF-mediated potentiation of DA transmission in the NAc was dependent on activation of either nAChRs or mAChRs.

Repeated exposure to stress or psychostimulant have been shown to alter the expression and/or function CRF peptide, CRF-R1 and CRF-R2 in a region dependent fashion (Curtis et al., 1995; Day et al., 1998b; Lemos et al., 2012; Liu et al., 2005; Reyes et al., 2006; Reyes et al., 2008; Waselus et al., 2009). We previously reported that repeated stress disrupts CRF’s ability to enhance dopamine in the NAc. This cellular alteration was accompanied by a switch in the behavioral response to CRF directly infused into the NAc (Lemos et al., 2012). In stress-naïve animals CRF produces a conditioned place preference suggesting that animals will bias their time allocation to contexts associated with endogenous CRF release specifically in the NAc. In contrast, animals subjected to repeated stressor exposure avoid contexts paired with intra-NAc infusion of CRF indicating a fundamental switch in the “perception” of CRF (Lemos et al., 2012). Therefore, we tested the effects of CRF on cholinergic activity in both stress-naïve and stress-exposed animals.

## Results

### CRF enhances CIN firing in DS and NAc

CINs make up only 1-2% of neurons within DS and NAc, which makes recording from these neurons in a high throughput fashion challenging. To aid with visualization of striatal CINs, recordings were carried out in *ChAT-ires-Cre*^+/−^ mice crossed with the Cre-reporter mouse line Ai14 tdTomato (CIN-reporter). We confirmed that red fluorescently-labeled Cre+ cells showed immunoreactivity for choline acetyltransferase (ChAT) and displayed the classic electrophysiological and morphological properties of CINs (i.e. spontaneously active, *I*_H_ sag, accommodation and large soma) (SF1). As seen in many other spontaneously active neurons, CINs showed a “run-down” of the spontaneous firing whole-cell configuration, due to the dialysis of the cytoplasm by the internal solution. For this reason, the majority of electrophysiological recordings were conducted in cell-attached configuration with gigaOhm seal maintained throughout the recording.

Bath application of CRF (100 nM) on to the brain slices significantly increased the spontaneous firing of CINs in both the NAc and DS (combined dorsolateral and dorsomedial) (NAc: 2.6 Hz to 6.2 Hz; DS: 0.7 Hz to 2.7 Hz, 2-way RM ANOVA, main effect of drug: F_1,19_ = 99.83, post-hoc Sidak’s t-tested, *ps* < 0.0001, n = 10-12, Figure 1a, b). When normalized to baseline, CRF increased CIN firing by 253 ± 22% and 507 ± 96% of baseline for the NAc and DS respectively (Figure 1c, d). Vehicle application (stock: 0.01% acetic acid in water, final: 0.00001% acetic acid in ACSF) produced a 23 ± 10% decrease in firing over time, likely due to run-down over time. CINs in the DS had a significantly lower frequency of spontaneous AP firing compared to CINs recorded in the NAc (2-way RM ANOVA, main effect of region: F_1,19_ = 30.29, *p* < 0.0001). Qualitatively, it appeared that cells with lower firing rates at baseline had larger CRF-mediated increases in firing. To formally assess this negative correlation, we plotted the magnitude of the CRF effect in percent as a function of the baseline firing rate in Hz on a semi-log plot (Figure 1e). This analysis yielded an R^2^ value of 0.51 suggesting that baseline firing rate accounts for approximately 50% of the size of CRF’s potentiation of CIN firing. Next, we assessed the concentration dependence of the CRF effect was assessed next in NAc CINs. For this, we screened cells with baseline firing rates between 1 and 5 Hz, which provided similar average baseline firing rates across the experiments with different CRF dose (Figure 1f). The CRF concentration dependence curve can be fitted to sigmoidal equation with EC50 of 8.7 nM, demonstrating the high sensitivity of this novel mechanism of CRF action in the striatum (Figure 1g,h).

**Figure 1.**
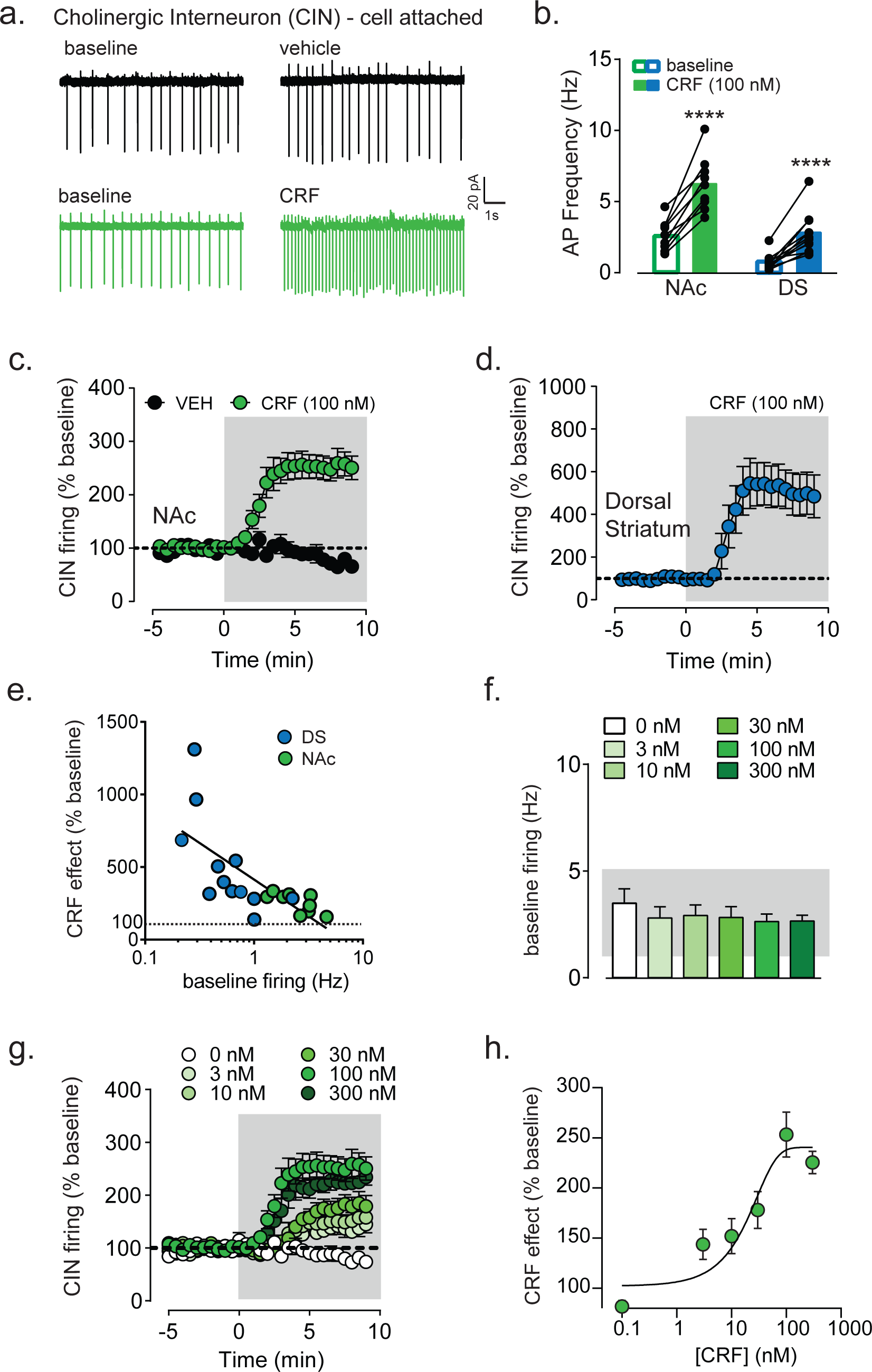
CRF increases the spontaneous firing of striatal CINs. a) Representative traces from CINs recorded in the NAc prior to and after vehicle (top) or CRF (100 nM, bottom) is bath applied. b) Firing rate prior to and following CRF application in CINs recorded in NAc or DS. c, d) Time course of CRF/vehicle effects on CIN firing frequency normalized to the baseline, recorded in NAc (c) or DS (d). e) Semi-log plot of CRF effect (% baseline) as a function of baseline firing frequency recorded from individual CINs in NAc (green) or DS (blue). f) Average baseline firing frequencies of NAc CINs from each [CRF] (0, 3, 10, 30, 100, 300 nM) group. g) Time course of CRF effects on normalized CIN firing frequency with different [CRF] (0, 3, 10, 30, 100, 300 nM). h) CRF effect (% baseline) was fit to a sigmoid curve as a function of CRF concentration.

### CRF-R1 receptors are required and are expressed in CINs

A combination of RNAscope *in situ* hybridization and pharmacology was used to identify the receptor subtype responsible for the CRF-mediated increase in CIN firing. RNAscope *in situ* hybridization was performed on thin brain sections (16 µm) containing the DS and NAc in which a probe against *ChAT* (CIN) mRNA was multiplexed against probes for either *crh1* (CRF-R1), *crh2* (CRF-R2) or *crh* (CRF) mRNA. Three to four sections from 3 animals were imaged using confocal microscopy and a combination of automated and manual analysis was performed using Fiji/ImageJ software. Cells showing at least five labeled puncta tightly clustered around the nucleus were considered positive for CRF-R1, R2 or CRF. The analysis showed that 92% and 90% of ChAT positive neurons contained CRF-R1 mRNA in the NAc and DS respectively (Figure 2a,b, SF2). In contrast, there was no evidence of co-localization of CRF-R2 or CRF mRNA with ChAT mRNA in either the NAc or DS (Figure 2a-d, SF2).

**Figure 2.**
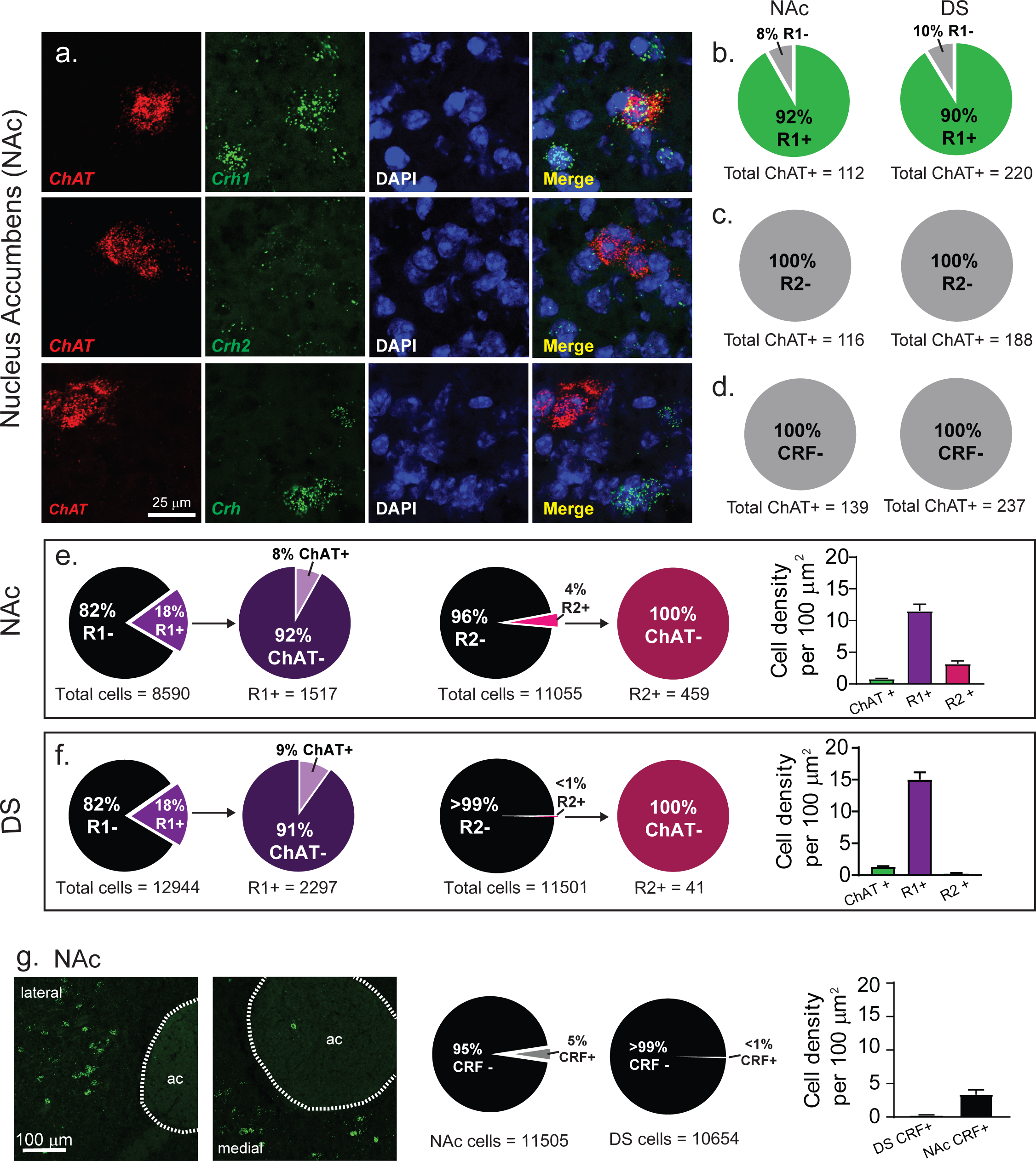
NAc and DS CINs ubiquitously express CRF-R1. a) Representative fluorescent *in situ* hybridization images of CIN marker *ChAT* mRNA (red), mRNAs (green) for *Crh1* (CRF-R1, top), *Crh2* (CRF-R2, middle), or *Crh* (CRF, bottom), and DAPI nuclear stain (blue) in NAc. b, c,d) Quantification of *ChAT* (CIN) co-localized with mRNAs for *Crh1*(b), *Crh2* (c), or *Crh* (d) in NAc (right) or DS (left). e) Quantification of DAPI cells co-localized with mRNAs for *Crh1* (left) and *Crh2*(middle) in the NAc. The number of CRF-R1+, CRF-R2+, and ChAT+ cells per 100 µm^2^ (right). f) Quantification of DAPI cells co-localized with mRNAs for *Crh1* (left) and *Crh2*(middle) in the DS. The number of CRF-R1+, CRF-R2+, and ChAT+ cells per 100 µm^2^ (right). g) Representative fluorescent images and quantification of DAPI cells colocalized wutg nRNA for *Crh* in the NAc or DS. The number of CRF+ cells per 100 µm^2^ in NAc or DS.

There is a substantial literature indicating that the CRF-R1s are expressed in cholinergic projection neurons of the basal forebrain (Sauvage and Steckler, 2001). As a control, we also quantified labeling in the basal forebrain of the same tissue sections. Analysis of ChAT+ neurons in the diagonal band area where most cholinergic neurons that project to the hippocampus are found (Ballinger et al., 2016), showed 45% of ChAT+ cells were also co-labeled for CRF-R1 mRNA while only 1% were co-labeled for CRF-R2 mRNA (SF2). Thus, to our surprise, expression of CRF-R1 is even more prominent and robust on CINs within the striatum than on cholinergic neurons in the basal forebrain.

We also observed CRF-R1 mRNA expression in cells that did not express mRNA for ChAT in both the DS and NAc. 18% of all DAPI labeled cells were positive for CRF-R1 receptors in both the DS and NAc (Figure 2e,f). The presence of cells expressing mRNA for CRF-R2 was negligible in the DS (<1%). However, in the NAc, 4% of cells were positive for mRNA for CRF-R2 (Figure 2e,f). Future work will be needed to determine the identity of those cells that are presumed noncholinergic neurons based on undetectable mRNA for ChAT and that are shown to express mRNA for CRF-R1 and R2.

With regard to the expression of mRNA for the neuropeptide itself, we found that 5% of cells in the NAc showed labeling for mRNA for CRF, but no cells were found in the DS (Figure 2g). In contrast, cells in PVN, BNST, and CeA showed robust expression of CRF mRNA using the same RNAscope probe (SF3). Notably, the lack of localization of mRNAs for CRF and ChAT is contrary to our previous finding of co-localization of immunostaining for CRF and ChAT (Lemos et al., 2012).

Given the strong evidence for the expression of CRF-R1 in CINs, we hypothesized that activation of CRF-R1 was required for the CRF-mediated increase in firing of CINs. For the remainder of our experiments, we focused our analysis on the neurons in the NAc and tested the ability of the selective antagonists for CRF-R1 (NBI 35695) and for CRF-R2 (Astressin 2B) to block the CRF effect. Slices were pre-incubated with either antagonist for at least 20 minutes and maintained in the antagonist for the duration of the experiment. In the presence of NBI 35695 (1 µM), CRF failed to increase CIN firing frequency. We also tested a different R1 antagonist, CP 154,526 (1 µM) which also reduced the effect of CRF, but did not block it completely. In contrast, Astressin 2B (300 nM) had no effect on the potentiation by CRF (295% increase from baseline) (NBI: 2.6 Hz to 2.2 Hz; Astressin 2B: 2.1 Hz to 4.8 Hz, 2-way RM ANOVA, interaction: F_1,11_ = 47.27, post-hoc Sidak’s t-test for AS2B, *p* < 0.0001, n = 6-7, Figure 3a-c). (DMSO vehicle vs. CP154.526, 2-way RM ANOVA, interaction: F27,243 = 5.714, *p* < 0.0001, n = 5-6, data not shown).

**Figure 3.**
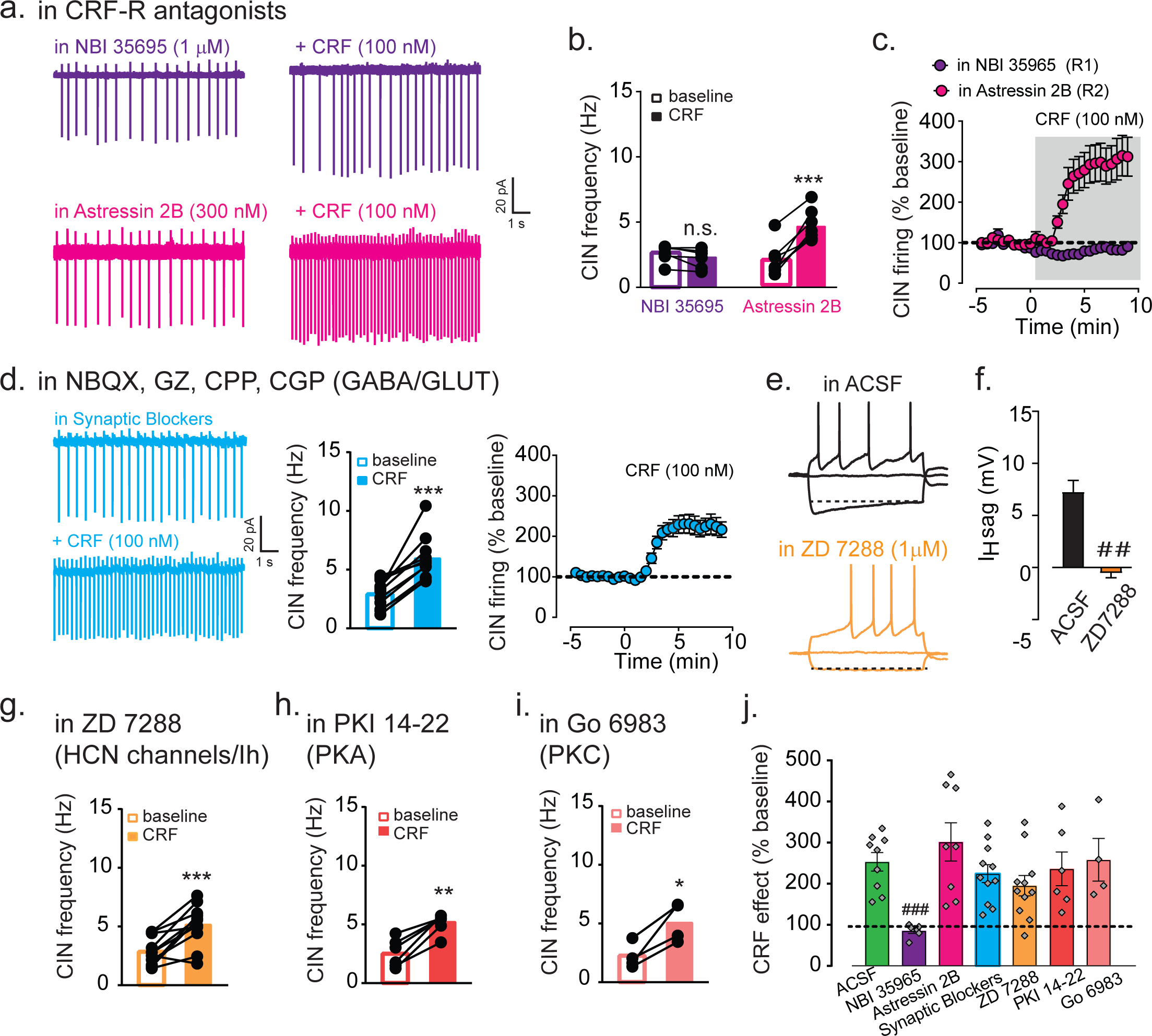
CRF mediated increase in CIN firing is driven by CRF-R1. a) Representative traces of CIN firing prior to and after CRF bath application in the presence of either NBI 35965 (CRF-R1 antagonist, 1 µM, top) or Astressin 2B (CRF-R2 antagonist, 300 nM, bottom). b) NAc CIN firing frequency prior to and after CRF application in each condition. c) Time course of CRF effect on CIN firing frequency normalized to baseline in each condition. d) (Left) Representative traces, (middle) CIN firing frequency, and (right) the time course of average CIN firing prior to and after CRF bath application in the presence of NBQX (20 µM), CPP (5 µM), CGP 55845 (2 µM), gabazine (5 µM). e) Representative voltage traces in response to different amount of current injection recorded from CINs in (top) ACSF or (bottom) ACSF + ZD 7288 (1 µM). f) Average *I*_h-_mediated “sag” in ACSF or ACSF + ZD 7288. g-i) CIN firing frequency prior to and following CRF application in the presence of ZD 7288 (g), PKI 14-22 (1 µM, h) or Go 6983 (3 µM, i). j) Summary of maximal CRF response (% baseline) for the experiments described in a-i.

### Evidence for CRF-mediated reduction of spike accommodation and dependence of cAMP activation

Pharmacological tools were used to investigate the mechanism underlying the increase in firing of action potentials using both cell-attached voltage-clamp and whole-cell current clamp recordings. The effect of CRF persisted in the presence of antagonists for GABA-A (gabazine, 5 µM), GABA-B (CGP 55845, 2 µM), NMDA (CPP, 5 µM), and AMPA (NBQX, 20 µM) receptors, indicating that the mechanism for CRF effects does not involve glutamate and GABA transmission and that the CRF effects is not through modulation of the release of these classical neurotransmitters onto to CINs (in the antagonists: 2.9 Hz to 5.9 Hz, paired t-test, *p* < 0.0001; 2-way RM ANOVA compared to in ACSF, F27,486 = 0.7891, *p* = 0.7680, n = 11, Figure 3d).

Previous work by Wanat and colleagues, showed that CRF enhances the tonic firing of VTA dopamine neurons by increasing *I*_h_ conductance through a PKC dependent mechanism (Wanat et al., 2008). Since striatal CINs show similar pacemaker properties as VTA dopamine neurons in an *ex vivo* slice preparation, it seemed plausible that CRF-R1s in CINs may be utilizing a similar mechanism. We first confirmed that preincubation with ZD 7288 (1 µM) can effectively block *I*_h_ conductance in CINs by measuring the hyperpolarization-induced “sag” in current clamp configuration. (*I*_h_ was reduced from 7.2 ± 0.1 mV to −0.5 ± 0.5 mV by ZD 7288; t-test, p < 0.0001, n = 11-12, Figure 3e-f). However, this same concentration of ZD resulted in only a modest reduction of the CRF effect under cell-attached configuration (firing from 2.9 ± 0.4 in baseline to 5.2 ± 0.5 Hz after CRF; paired t-test, *p* = 0.0007, n = 11, Figure 3g). It is worth mentioning that there was a significant difference in the time course of the CRF effect (2-way RM ANOVA, interaction, F25,450 = 2.6, *p* < 0.0001, n = 9-11, data not shown).

The CRF effect on CIN firing frequency was unaffected in the presence of PKA (PKI 14-22, 1 µM) or PKC (Go 6983, 3 µM) inhibitors (PKI 14-22: 2.5 ± 0.5 to 5.1 ± 0.8 Hz, paired t-test, *p* == 0.002; 2-way RM ANOVA in comparison to ACSF, interaction, F27,351 = 1.17, *p* = 0.26, n = 6, Figure 3e; Go 6983: 2.2 ± 0.5 to 5.1 ± 0.8, paired t-test, *p* = 0.02; 2-way RM ANOVA in comparison to ASCF, interaction, F27,297 = 1.8, *p* = 0.01, n = 4, Figure 3h-i). We again observed that the time course to achieve maximal CRF effect was delayed in the presence of Go 6983. Overall, when assessing the effect of all pharmacological blockers on the CRF effect expressed as percent of baseline firing, only the CRF-R1 antagonist NBI 35695 showed a significantly effect (1-way ANOVA, Dunnett’s post-hoc t-test, p = 0.003, Figure 3j).

Current clamp recordings were used to further challenge the lack of effects by some blockers and for testing inhibitors that were impermeable to the cell membrane and required introduction via the recording pipette. Within 5 min after achieving whole-cell current clamp recordings, cells often became hyperpolarized and eventually stopped firing. Cells were maintained at ∼ −70 mV throughout the experiment (SF1) and current steps (800 ms duration) were applied from these resting potentials. This protocol enabled us to analyze the effect of CRF on different components of CIN excitability including *I*_h_ sag, membrane resistance accommodation, and depolarization induced block of AP firing (Figure 4, ST1). As shown in Figure 4a, CINs showed the canonical spike accommodation in response to the current steps. AP frequency from the first 400 ms (FIRST) was higher than form the last 400 ms (LAST), which was true when measured under all conditions: ACSF, VEH and CRF (2-way RM ANOVAs, interactions: baseline/VEH, ps < 0.0001; baseline/CRF, ps < 0.0001, Figure 4 a-f).

**Figure 4.**
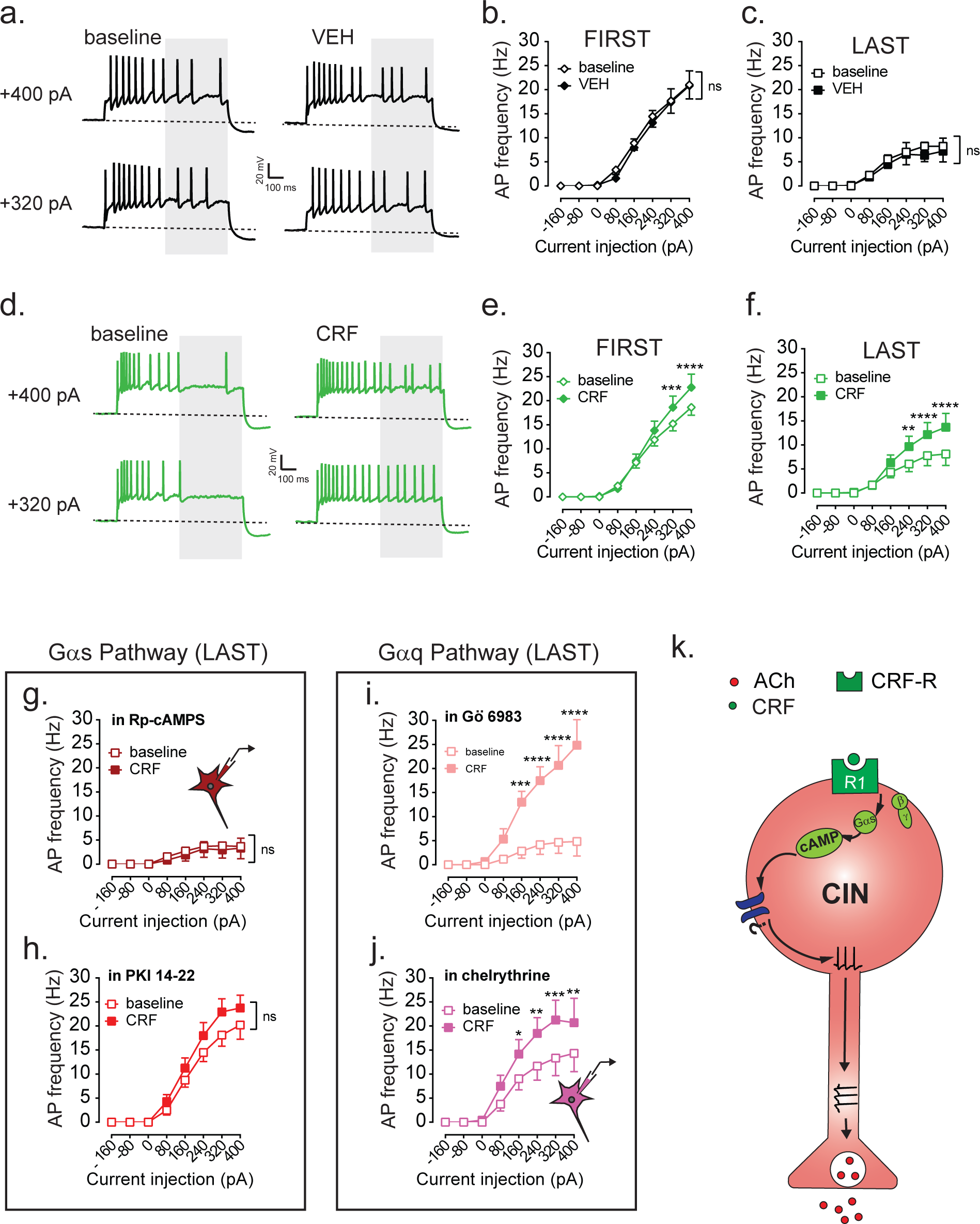
CRF mediated increase in CIN firing requires cAMP. a) Representative voltage traces recorded from striatal CIN in response to depolarizing current injections (+320, +400 pA) before and after vehicle application. b) Average input-output response of NAc CINs for the first 400 msec of an 800 msec sweep before and after vehicle application. c) Average input-output response of NAc CINs for the last 400 msec of an 800 msec sweep before and after vehicle application. d) Representative striatal CIN voltage response to depolarizing current injections (+320, +400 pA) before and after CRF application. e) Average input-output response of NAc CINs for the first 400 msec of an 800 msec sweep before and after CRF application. f) Average input-output response of NAc CINs for the last 400 msec of an 800 msec sweep before and after CRF application. g-j) Average input-output curve (last 400 msec) for CINs recorded prior to and following CRF (100 nM) bath application in cells with Rp-cAMPS present in the internal solution (g), slices incubated and maintained in PKI 14-22 (h), cells with chelrythrine present in the internal solution (i) and in slices incubated and maintained in Go 6983 (j). k) Summary cartoon detailing the mechanism by which CRF increases CIN firing.

With the vehicle application, there was no significant change in AP frequency from the whole period, the first half, or the last half (2-way RM ANOVA, ps = 0.99, 0.99, 0.98, respectively, n = 9, Figure 4a-c). In contrast, CRF significantly increased AP frequency from the whole period, the first half, and the last half (2-way RM ANOVA, ps < 0.0001, n = 12, Figure 4d-f). The increase of AP frequency by CRF was most prominent during the last 400 ms. The inverse of the slope (or 1/slope) from the input-output curves is indicative of the magnitude of injected current needed to generate each action potential. The 1/slope was not significantly affected by the vehicle (from 18 to 17 pA/spike for the first half; from 54 to 64 pA/spike. for the last half; Figure 4b,c). In contrast, CRF decreased the 1/slope for the last half selectively from 49 to 27 pA/spike (first half: from 20 and 15 after CRF; Figure 4h,i). The steeper slope in the presence of CRF, may be evidence of CRF actions relieving the depolarization induced block of AP frequency, which is more prominent during the last 400 ms. Consistent with findings from cell-attached recordings, incubation in the *I*_h_ inhibitor ZD 7288 only had a modest effect on the robustness and reliability of the CRF effect on AP frequency, particularly when analyzing the entire sweep or first 400 ms (SF4) This indicates that *I*_h_ current is not required for this effect.

CRF-R1 activation engages the Gαs signaling pathway in other cell types (Hauger et al., 2006). Recruitment of Gαs can initiate cAMP-dependent recruitment of both protein kinase A and other effectors such as guanine nucleotide exchange factors (EPACs). We used a cAMP analogue Rp-cAMPs (100 µM) was included in the internal solution and allowed to dialyze into the cell; this treatment blocks endogenous cAMP signaling. Under these conditions, CRF failed to increase the AP frequency (2-way RM ANOVA, interaction: p’s = 0.9778, 0.7804, 0.9857; main effect of CRF: p’s = 0.6307, 0.4869, 0.6159 for the full 800 ms, the first 400 ms, the last 400 ms, respectively; n = 7, Figure 4g, SF4). When we directly inhibited PKA with PKI 14-22, CRF still significantly increased the AP frequency during the first 400 ms (2-way RM ANOVA, interaction: ps = 0.0001, 0.0002 respectively, n = 8, Figure 4h, SF4), but failed to increase during thelast 400 ms (2-way RM ANOVA, interaction: p = 0.2527; main effect of CRF: p = 0.0732, n = 8, Figure 4n, SF4). These data indicate that while activation of PKA may contribute modestly to CRF’s ability to increase CIN firing, cAMP activation is required for the CRF effect on CINs.

We also tested two different PKC inhibitors, chelrythrine (3µM) in the internal solution or Go 6983 (1 µM) applied to the bath. CRF robustly enhanced AP frequency over a range of depolarizing currents regardless of how we looked at the sweep in the presence of chelrythrine or Go 6983 (2-way RM ANOVA, interaction: ps = 0.0025 and 0.0001; main effect of CRF: ps = 0.02, 0.009 for chelrythrine and Go 6983 respectively with *, **, ***, **** signify Sidak’s post-hoc t-tests with p values < 0.05, 0.01, 0.001, 0.0001 respectively, n = 5-6,Figure 4i, j, SF4).

Taken together, these data indicate that CRF enhances CIN firing through multiple effectors, all of which rely on cAMP. The effect of CRF seems to be particularly prominent when measuring the AP frequency when the depolarization induced block common to CINs is most apparent. PKA and *I*_h_ may play a modest role and may be necessary in order for CRF to achieve and its typical reliable and maximal effect size, however it is not required for the effect to occur. Based on both the cell-attached voltage clamp and current clamp experiments, it does not appear that the CRF-mediated increase in CIN firing frequency is reliant on PKC activation (Figure 4k).

### CRF potentiation of DA transmission does not require nicotinic acetylcholine receptors

Given the profound effect of CRF on CIN firing frequency, we hypothesized that the potentiation of DA transmission by CRF is mediated by activation of acetylcholine receptors on the axons of DA neurons. Thus, we predicted that blockade of nicotinic receptors with the β2 nAChR antagonist DHβE would block CRF potentiation of electrically evoked DA transient (eDA). Using fast scan cyclic voltammetry (FSCV) in the same *ex vivo* slice preparation, we replicated the main finding that CRF increases the peak dopamine transient by 21 ± 4% above baseline (one-sample t-test vs. 0, p = 0.003, n = 16, Figure 5a, SF5). More importantly, CRF potentiation was significantly greater than vehicle control (2-way RM ANOVA, interaction of CRF vs. VEH, F_13,338_ = 5.014, p < 0.0001, n = 12-16, Figure 5a).

**Figure 5.**
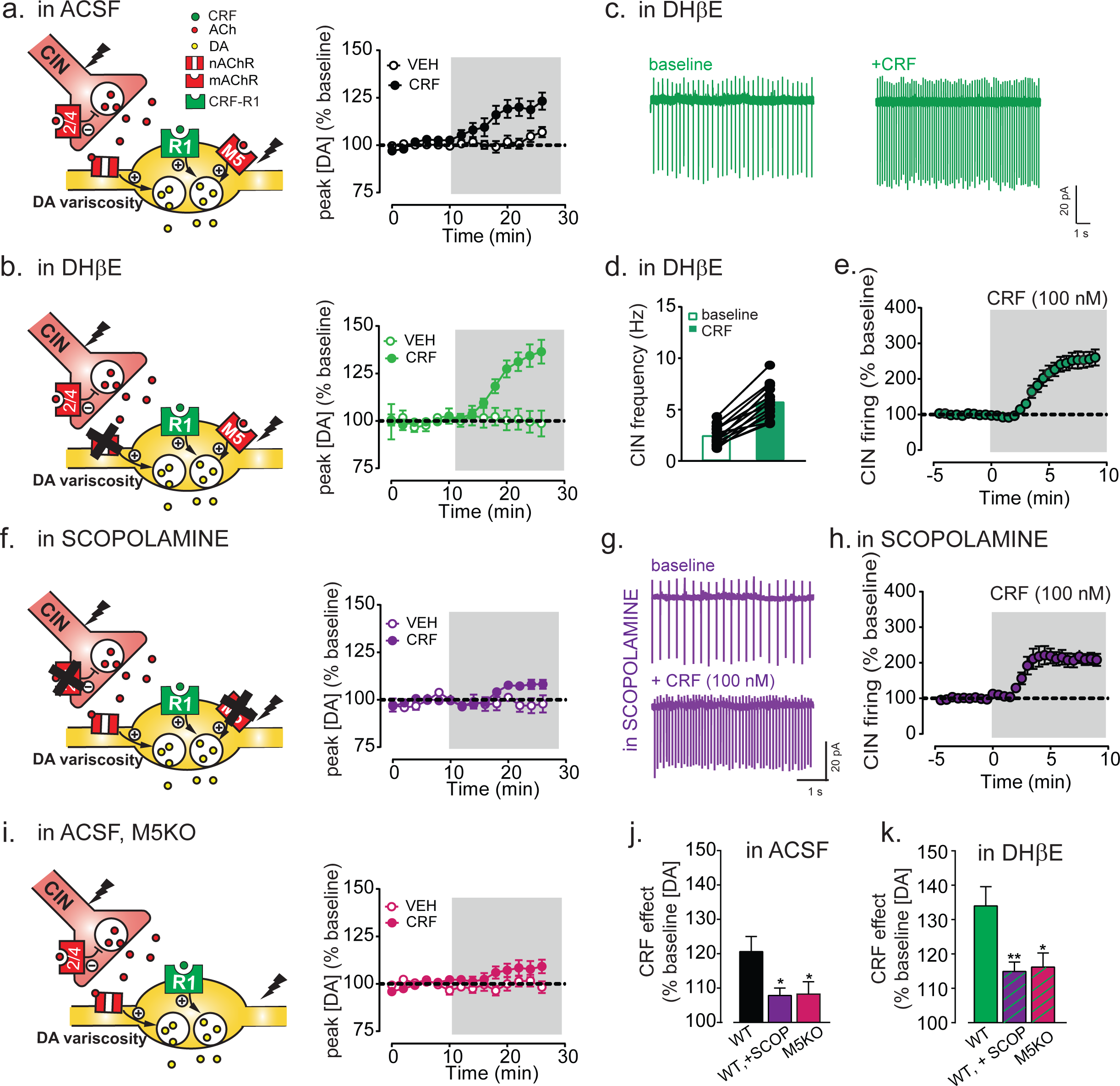
CRF potentiation of DA transmission requires mACh-R type 5 transmission. a) Schematic and mean time course of electrically evoked DA transients recorded before or after CRF or vehicle application in WT control NAc slices incubated and maintained in ACSF. b) Schematic and mean time course of electrically evoked DA transients recorded before or after CRF or vehicle application in WT control (in-house C57Bl6/J) NAc slices incubated and maintained in ACSF + DhβE. c-e) Representative traces (c), firing rate (d) and normalized timecourse (e) of NAc CINs prior to and following CRF application in slices incubated and maintained DhβE. f) Schematic and mean time course of electrically evoked DA transients recorded before or after CRF or vehicle application in WT control NAc slices incubated and maintained in ACSF + Scopolamine. g) Schematic and mean time course of electrically evoked DA transients recorded before or after CRF or vehicle application in WT control NAc slices incubated and maintained in ACSF + DhβE Scopolamine. h) Schematic and mean time course of electrically evoked DA transients recorded before or after CRF or vehicle application in M5KO NAc slices incubated and maintained in ACSF. i) Schematic and mean time course of electrically evoked DA transients recorded before or after CRF or vehicle application in M5KO (C57Bl6/J back-crossed) NAc slices incubated and maintained in ACSF+ DhβE. j) Average maximal CRF effect on peak [DA] (% baseline) in experiments conducted in ACSF (a, f,h). j) Average maximal CRF effect on peak [DA] (% baseline) in experiments conducted in ACSF + DhβE (b, g,i). l, m) Representative traces (l) and normalized time course (m) of NAc CINs prior to and following CRF application in slices incubated and maintained scopolamine.

Application of DHβE (1 µM) alone reduced the amplitude of eDA by more than 50% of the baseline, isolating the component of DA transient evoked by direct electrical stimulation of DA axon fibers in the striatum (Zhang et al., 2002; Shin et al., 2015). However, CRF still potentiated eDA by 34 ± 5.6% in the presence of DHβE (one-sample t-test vs. 0, p < 0.0001, n = 16, 2-way RM ANOVA vs. VEH, interaction: F_13,273_ = 7.082, p < 0.0001, n = 7-16, Figure 5b, SF5). This potentiation by CRF in DHβE was modestly enhanced compared to ACSF at some time points although this was not significant when assessing the collapsed maximal effect (unpaired t-test, p = 0.07, SF5). When we probed this further using electrical stimulation of a train of 5 pulses at 10 Hz, a stimulation closely resembling *in vivo* phasic DA transmission, the potentiation by CRF was significantly enhanced in DHβE compared to ACSF (SF5, Supplementary Results & Discussion). As a control, we demonstrated that incubation in DHβE did not alter the maximal CRF mediated increase in CIN firing frequency, though the time to maximal effect was slightly delayed (in DHβE: baseline: 2.45 ± 0.26 Hz, CRF: 5.71 ± 0.46 Hz, p < 0.0001; 2-way RM ANOVA compared to in ACSF, F_27,567_ = 2.62, p < 0.0001, n = 14, Figure 5c-e).

We have previously shown that CRF-mediated potentiation of DA transients is blocked by pre-treatment with the CRF-R1 antagonist (Lemos et al., 2012). Here we then tested whether this same CRF-R1 antagonist could reverse the potentiation after CRF application. Interestingly, we found that the CRF-R1 antagonist CP 154,526 (1 µM) could not reverse the CRF-mediated potentiation of eDA, even after an hour of CRF washout and CRF-R1 antagonist chase (SF6). This data suggests that CRF may initiate a long-lasting change in the striatal circuitry that triggers a long-term potentiation-like effect on DA transmission.

### CRF potentiation of dopamine transmission is dependent on M5 receptor activation

We and others have previously shown that muscarinic M5 receptors potentiate dopamine transmission, particularly when triggered by activation of dopamine fibers optogenetically or by pharmacological isolation (Schmidt et al., 2010; Shin et al., 2015; Zhang et al., 2002). In this study, we replicated these findings (SF7, see Supplementary Results & Discussion). We then tested whether the activation of mAChR is required for CRF-mediated potentiation of DA transients. Scopolamine (1 µM), a non-selective mAChR antagonist, reduced the CRF-mediated potentiation by 50%, both in control ACSF and in DHβE conditions (in ACSF + SCOP, CRF vs. VEH: 2-way RM ANOVA, interaction: F_13,386_ = 2.158, p = 0.01, n = 6-16, Figure 5f, j; in DHβE+SCOP: 2-way RM ANOVA, interaction: F_13,281_ = 4.567, p < 0.0001, n = 11-13, Figure 5k). While smaller, CRF still showed a significant potentiation of eDA compared to vehicle. These data suggest that activation of muscarinic receptors mediates a large part, but not all of, the CRF potentiation of DA transmission.

In mice with constitutive genetic deletion of M5 mAChR (M5KO), CRF enhanced dopamine transient amplitude compared to vehicle application but did so less than in slices from control mice and to a similar degree as seen in the presence of the non-selective muscarinic receptor blocker scopolamine (control = 20.6 ± 4.4%, M5KO= 8.25 ± 2.6%; SCOP= 7.85 ± 2.2%, 1-way ANOVA, F = 4.49, p = 0.017, post-hoc t-test vs. control: ps = 0.019, 0.035 for SCOP or M5KO respectively, post-hoc t-test SCOP vs. M5KO, p = 0.9998, n = 13-16, Figure 5i-j).

Similar results were obtained when CRF was tested on DA transients in the presence of DHβE (in DHβE: control= 34.0 ± 5.6%, M5KO= 16.2 ± 2.1%, SCOP= 14.9 ± 2.7%, 1-way ANOVA, F = 5.55, p = 0.008, post-hoc t-test vs. control: ps = 0.009, 0.037 for SCOP or M5KO respectively, post-hoc t-test SCOP vs. M5KO, p = 0.9846, n = 8-16, Figure 5k).

Altogether, these findings provide evidence for the existence of two independent mechanisms for CRF-mediated potentiation of DA transmission, one that requires activation of M5 mAChR receptors and a small one independent of muscarinic receptor activation that is likely mediated by CRF receptors on dopamine terminals, based on anatomical data (Lemos et al., 2012). We tested whether CRF effect on CIN firing frequency also required muscarinic receptor activation and found that CRF still increase CIN firing in the presence of scopolamine (from 2.8 ± 0.35 Hz at baseline to 5.3 ± 0.38 after CRF, p < 0.0001, n = 11, Figure 5g,h), and to a comparable degree to the CRF effect in ACSF (2-way ANOVA, interaction, p = 0.2181). Thus, these data indicate that the requirement for muscarinic receptors is downstream of the CRF actions on CIN firing and it suggests that the M5 mAChR receptors that contribute to the CRF potentiation of DA transmission are localized to DA axon projections and/or terminals, in agreement with previous work (Shin et al., 2015).

### Repeated stress differentially affects CRF function and mRNA expression based on cell type

Stressor exposure has been shown to alter the mRNA expression, protein distribution and function of CRF and its two receptors (Milan-Lobo et al., 2009; Reyes et al., 2008; Waselus et al., 2009). We conducted a series of studies where we probed CRF function as well as mRNA expression of *Crh* and *Crh1* in animals exposed to repeated stress (FSS) and compared them to naïve mice kept in homecage (Figure 6a-c). Mice with fluorescently labeled CINs (ChAT-ires-Cre^+/−^; Ai14 tdTomato) were exposed to two days of repeated swim stress and studied 7 days after last stressor exposure (Figure 6b). Slices containing the NAc were prepared and half were used for DA transient measures using FSCV and the other half were used for cell-attached electrophysiology recordings from CINs, providing an internal control for CRF function on the two effectors (Figure 7b). The results of the FSCV experiments replicated previous findings showing that exposure to repeated swim stress caused a persistent reduction of CRF-mediated potentiation of eDA. The maximal CRF effect was 20.5 ± 3.6 % increase in naïve mice while there was only a 11.6 ± 2.5% increase in animals that underwent FSS (unpaired t-test, p = 0.048, n = 7-11; Figure 6d,e). We hypothesize that the reduction could be due to selective loss of one of the two independent mechanisms of CRF action: the M5-dependent or the M5-independent mechanism. We tested the direct effect of a muscarinic agonist oxotremorine (Oxo-M) on slices from naïve and FSS animals in the presence of DHβE to avoid cross talk between ACh receptors. In naïve animals, application of Oxo-M produced an increase of 14 ± 4%. in the steady state amplitude of eDA transient. Subsequent application of CRF further increased eDA to 29.1 ± 4.6% from initial baseline, revealing an M5-independent component of around 15% (Figure 6f), in agreement with data from Figure 5 showing approximately 50:50 contribution of each mechanism. In slices from FSS animals, Oxo-M still produced an increase in eDA transient amplitude with similar early rise of 24 ± 3% (compared to 22 ± 2% in naïve) and a steady state increase of 10 ± 4.5 % (compared to 14% in naïve; Figure 6f). There was no statistical difference between the groups (2-way RM ANOVA, post-hoc t-test, p = 0.99). However, in FSS group, subsequent application of CRF failed to further increase eDA amplitude (4 ± 5 % in FSS compared to 16 ± 5% in naïve from amplitude in Oxo-M, n = 8-10;2-way RM ANOVA of FSS vs. naïve, interaction: F_22,352_ = 2.65, p = 0.0001, post-hoc t-test, p = 0.02, Figure 6f). These results strongly suggest that exposure to repeated stress selectively impairs the M5-independent component of CRF action while sparing the M5 dependent component intact.

**Figure 6.**
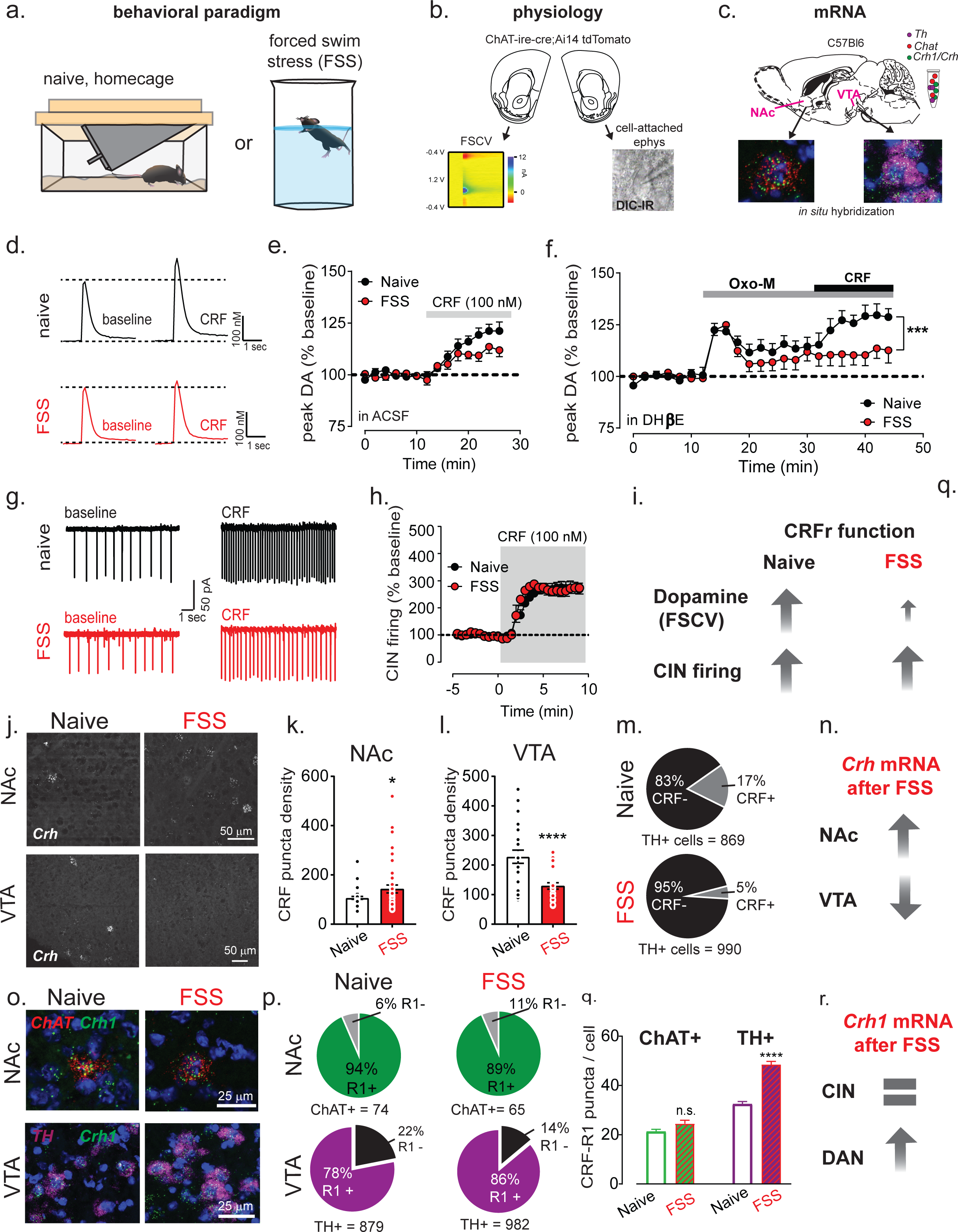
Repeated stress differentially effects CRF-R1 function and expression in a cell-type dependent fashion. a) Schematic depicting stress-naïve vs. stressor-exposed behavioral manipulation and comparison. b) Schematic depicting within animal acquisition of both CIN firing frequency and fscv data. c) Schematic depicting within animal acquisition *in situ* hybridization data for *ChAT, Crh1/Crh* and *Th* expression. d) Representative traces of electrically evoked DA transients prior to and following CRF application in NAc slices from stress-naïve or stressor exposed mice. e) Mean peak amplitude of electrically evoked DA transient over time in response to CRF application normalized to baseline in NAc slices from stress-naïve and stressor-exposed mice. f) Mean peak amplitude of electrically evoked DA transients over time in response to OxoM followed by Oxo-M + CRF application normalized to baseline in NAc slices from stress-naïve and stressor-exposed mice. g) Representative traces of CIN firing frequency prior to and following CRF application in slices taken from either stress-naïve or stressor-exposed *ChAT-ires-Cre^+/−^;Ai14*. h) CIN firing frequency over time in response to CRF application normalized to baseline for slices from stress-naïve or stress-exposed mice. i. Summary cartoon of CRF function of CIN firing and DA transmission in naïve and stress-exposed mice.j) Representative fluorescent *in situ* hybridization images of *Crh* (CRF) in NAc (top) or VTA (bottom) of the same stress-naïve and stress-exposed mice. k) Mean # of CRF mRNA puncta in the NAc in naïve vs. stress-exposed mice. l) Mean # of CRF mRNA puncta in the VTA of naïve vs. stress-exposed mice. m) Quantification of TH+/CRF+ cells in the VTA in naïve and stress-exposed mice. n) Summary cartoon of stress-induced alterations in CRF mRNA in NAc and VTA. n) Representative fluorescent *in situ* hybridization images of *Crh1* (CRF-R1) and *ChAT* (CINs) in NAc (top) or *Th* (DAN) and *Crh1* (CRF-R1) in VTA (bottom) of the same stress-naïve and stress-exposed mice. o) Quantification of ChAT+/CRF-R1+ cells in the NAc (top) or TH+/CRF-R1+ cells in the VTA (bottom) in naïve and stress-exposed mice. p) Mean # of CRF-R1 mRNA puncta per cell in naïve vs. stress-exposed mice. q) Summary cartoon of stress-induced alterations in CRF-R1 mRNA in NAc and VTA.

In agreement with this interpretation, we found that prior stressor exposure did not alter the CRF-mediated increase in CIN firing (267 ±19% in FSS vs 273 ± 22% in naïve, n = 8-14; unpaired t-test, p = 0.8637, 2-way RM ANOVA no interaction: F_27,540_ = 0.7259, p = 0.8433, Figure 6g,h). Thus, altogether the findings suggest no changes to the function/signaling of CRF-R1 receptors on CINs nor M5 receptor on dopamine terminals. These led us to speculate that stress exposure induces changes in the function and/or expression of CRF-R1 receptors expressed on DA neurons and/or terminals, which we inferred mediate the M5-independent mechanisms of CRF action. Possible changes in receptor expression were assessed using RNAscope method using probes for the CRF-R1 mRNA, *Crh1,* in both the NAc and VTA. We also assessed the expression of the CRF peptide itself using primers for *Crh* mRNA.

Figure 6j-n summarize the results of the *in situ* hybridization experiments for *Crh* mRNA in the NAc and VTA. We found increased density of *Crh* mRNA puncta in the NAc of FSS animals compared to naive (143 ± 16 in FSS vs 107 ± 6 puncta/425 µm^2^ in naive, t-test, p = 0.03, n = 44-46 unique images from 3 animals each, Figure 6j,k,n). Surprisingly, there was a decrease in the density of *Crh* mRNA puncta in the VTA of FSS animals. The decrease was observed when quantifying both the overall VTA area (130 ± 11 in FSS vs 228 ± 22 in naïve, t-test, p = 0.0003, n = 22-23 images/3 mice each) and when quantifying puncta selectively on TH+ cells (4.8 ± 1 % in FSS vs 17 ± 2.8% in naïve, t-test, p = 0.0002, n = 879, 992 cells/3 animals each; Figure 6j-n). Note that, in agreement with a previous report (Grieder et al., 2014), we found that 17% of TH+ neurons were also positive for *Crh* mRNA.

For *Crh1* mRNA coding for CRF-R1 receptor, we quantified its expression in TH+ cells within the VTA and in ChAT+ CINs in the NAc using a multiplexing approach in which the three hybridization probes (*ChAT, TH and Crh1*) were tested in the same sagittal brain section in order to more reliably compare differences in expression across brain areas. In the VTA, contrast to our hypothesis, exposure to prior stressor significantly increased both the number of CRF R1+/TH + cells (naïve: from 78% in naïve to 86% in FSS, p = 0.03, Figure 7o-r) and the number of puncta per TH+ cell (naïve= 32 ± 1 puncta; FSS= −48.4 ± 1.3, n= 160-160 TH+ cells, p < 0.0001, Figure 7p,q). In the NAc, there was no differences in the number of *Crh1 mRNA* puncta per CIN cell (naïve= 21 ± 1 puncta; FSS = 25 ± 1.5, n= 67-71 *ChAT* + cells, p = 0.0589, Figure 6o-r).

**Figure 7.**
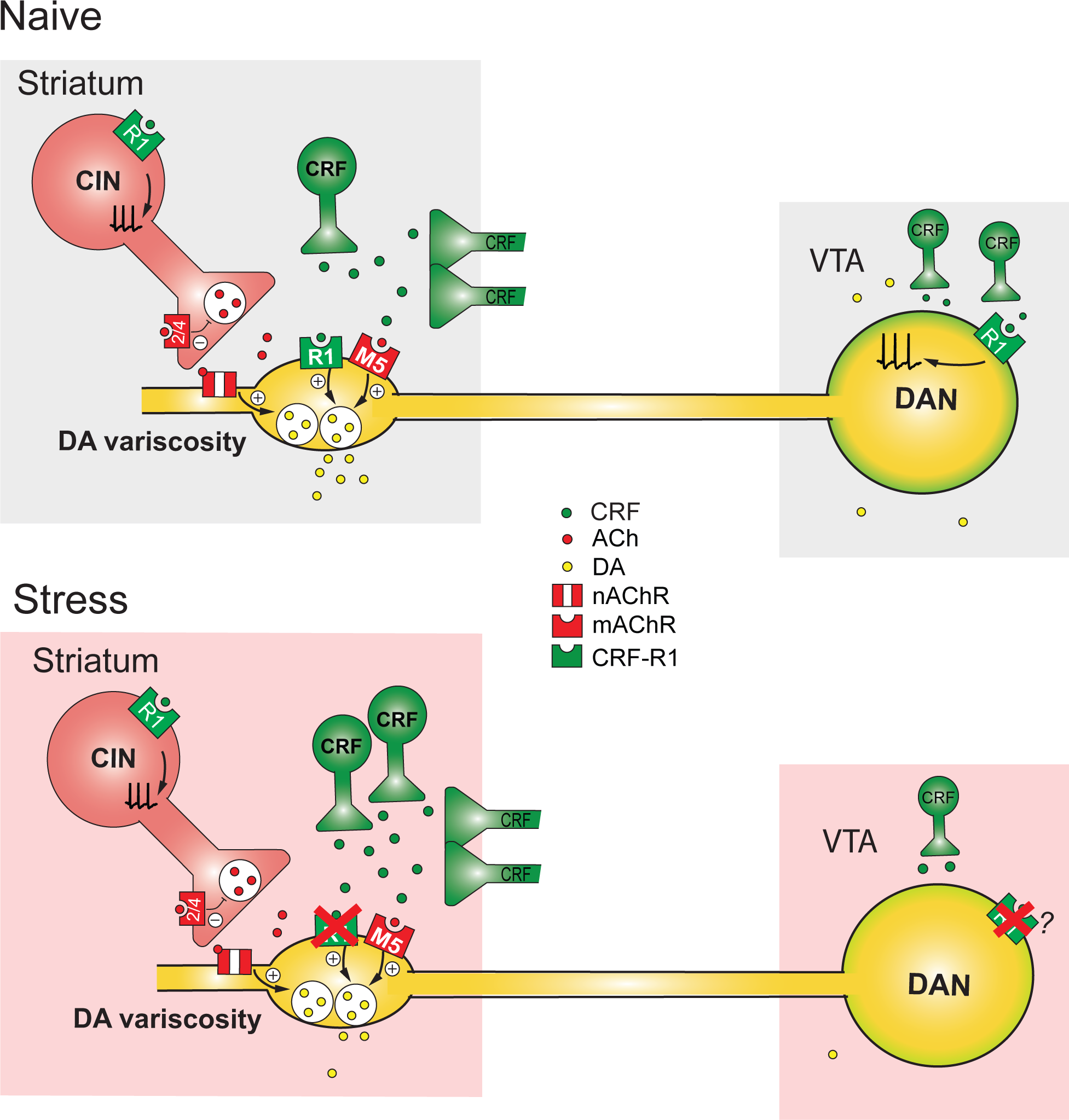
Summary cartoon of CRF function in the mesolimbic pathway in stress-naïve and stress-exposed mice. In stress-naïve mice, CRF increases CIN firing via activation of CRF-R1. CRF potentiates dopamine via direct activation of CRF-R1s on DA terminals and indirect activation of M5 receptors on dopamine terminals. This model also incorporates prior data from Wanat et al. (2008) and Beckstead et al. (2009) that showed that CRF-R1 increase DAN tonic firing and DA release in the VTA. CRF-containing cells are present in both the NAc and VTA. In stress-exposed mice, CRF potentiation of CIN firing is retained. CRF potentiation of dopamine transmission is reduced, likely due to disruption of CRF-R1s on dopamine terminals. This model also incorporates prior data from Beckstead et al. (2009) that showed restraint stress ablated CRF-R1 mediated release of DA in the VTA. Note, this was not shown using our repeated swim stress paradigm. Following stress, there are more CRF+ cells in the NAc and fewer CRF+ cells in the VTA.

Thus, we find exposure to repeated stress lowers the CRF-mediated potentiation of DA transmission in the accumbens by selectively affecting the M5-independent mechanism of CRF actions. As far as can be determined by our experiment, stress spares the CRF effect on CIN firing, and also the downstream potentiation of dopamine transmission mediated by muscarinic M5 receptors. Stress then produces selective alterations to CRF targets, even within the same region on the brain. It has been shown that CRF-R1 receptor can internalized upon agonist exposure (Reyes et al., 2006) and thus we speculated that prolonged CRF release during repeated stress could also trigger receptor internalization. We hypothesized that there may be a disruption of CRF receptor function due to increased CRF release in the NAc. While it is very difficult to assess CRF release directly, we were able to assess changes in CRF mRNA expression following repeated swim stress. We posited, based on previous anatomical work (Lemos et al. 2012), that the M5-independent component is mediated via CRF receptors localized to DA terminals. An additional explanation for the absence of the M5-independent component in the FSS group is through downregulation of CRF R1 function or expression at DA terminals.

## Discussion

### Novel CRF effector within the striatum: cholinergic interneurons

This study identifies a novel CRF effector within the dorsal and ventral striatum. A few previous reports showed sparse expression of CRF receptors in the striatum, (Henckens et al., 2016; Justice et al., 2008; Kuhne et al., 2012; Van Pett et al., 2000) but the cellular identity of this expression had not been addressed. Here we report that CRF-R1 receptors are expressed on CINs in the DS and NAc and that CRF robustly and reliably increases the spontaneous firing of action potentials in these neurons. The effect of CRF on CINs requires cAMP production and could involve suppression of spike accommodation. We propose that this novel effect of CRF might represent the underlying mechanism for a previously published effect of CRF in the NAc where elevation of ACh extracellular levels was seen in response to CRF (Chen et al. (2012)). In addition, our study provides evidence that this novel CRF action on CINs contributes to the CRF-mediated potentiation of DA transmission in the striatum.

### CRF indirectly activates ACh muscarinic M5 receptors to potentiate dopamine transmission

Our findings indicate that CRF potentiates dopamine transmission via two independent mechanisms: a) one that requires activation of ACh muscarinic M5 receptors and b) an M5-independent mechanism. Both these mechanisms required activation of CRF-R1 receptors, but we speculate that each requires the activation of CRF-R1 receptors expressed on different cell-types. Based on the *in situ* hybridization findings showing expression of *crh1* (the coding for CRFR1 receptors) in the CINs and the pharmacological data, we speculate that activation of CRF-R1 receptors on CINs causes the increase in spontaneous firing that increases the extracellular levels of Ach which in turn activate muscarinic M5 receptors on DA neuron axon terminals to potentiate DA transmission (Figure 7). Our model also posit that the M5-independent mechanism is mediated via CRF-R1 receptors on dopamine terminals that could directly potentiate DA transmission. CRF-R1 receptors are localized to DA fibers based on both immunohistochemistry and electron microscopy data (Lemos et al., 2012).

### CRF increases CINs firing and facilitates cholinergic transmission in the striatum

Acetylcholine release from the basal forebrain versus the striatum supports different, and in some cases opposite behaviors. Both stress and centrally (i.c.v.) administered CRF stimulate acetylcholine release in the hippocampus via activation of CRF-R1 receptors on cholinergic basal forebrain diagonal band projection neurons (Day et al., 1998a; Day et al., 1998b; Fernandes et al., 2018; Sauvage and Steckler, 2001). Moreover, this enhanced cholinergic transmission and nicotinic receptor activation in the hippocampus can lead to an increase in anxiety-and depression-like behavior (Fernandes et al., 2018; Mineur et al., 2013). Acetylcholine transmission in the striatum subserves a diverse set of behavioral responses that are dependent on the striatal subregion (i.e. NAc, DLS, DMS) and context in which ACh levels or CIN firing is being assessed. There is evidence that enhanced cholinergic signaling encodes salience regardless of valence and reflects reward prediction errors by shifting from a tonic to burst/pause mode (Deffains and Bergman, 2015). It has also been shown that modulation of CIN firing can support both approach or withdrawal behavior (Grasing, 2016; Hoebel et al., 2007), is important for reinforcement learning and for cognitive and behavioral flexibility (Aoki et al., 2018; Prado et al., 2017; Ragozzino, 2003). Here we show that CRF enhances CIN firing frequency regardless of region. Thus, we propose CRF acts to potentiate striatal ACh-dependent behaviors and that the specific behavior to be facilitated will depend on the striatal subregion where CRF is released. For example, we present evidence confirming that CRF peptide is present in a significant percentage of NAc neurons, but not DS neurons (Figure 2) and also (Merchenthaler, 1984; Peng et al., 2017). Thus, in the context in which CRF is released from these NAc CRF cells, we would predict that CRF would potentiate appropriate NAc-dependent ACh-dependent responses. This prediction should be directly tested in future studies.

CRF receptors were shown to couple to different G proteins depending on cell-type and agonist concentration. Here we showed that the CRF mediated increase in CIN firing requires cAMP production. Yet there was only a nominal effect of PKA or PKC inhibition on CRF’s ability to increase firing frequency. It is possible that CRF is exerting its effect on CIN firing through a cAMP dependent, but PKA independent mechanism. Indeed, there have been recent reports that CRF receptors can couple to EPAC2 (a guanine nucleotide exchange factor for the Ras-like small GTPases) in a cAMP dependent fashion (Smani et al., 2010; Traver et al., 2006; Van Kolen et al., 2010). EPACs have been shown to modulate the function of several different ion channel types including L-type calcium channels, AMPA receptors and voltage-gated potassium channels (Liu et al., 2016; Roscioni et al., 2008; Smani et al., 2010; Stott et al., 2016). Thus, it is plausible that CRF-R1 is stimulating EPACs to effect ion channel conductance and in turn firing frequency at one of these effectors.

### Repeated stress exposure selectively alters ACh-independent mechanisms and the expression of CRF receptor in the DA neurons but not CINs

Stress exposure produces plasticity in the CRF system that includes changes in peptide expression, effector coupling and function. CRF modulates the activity of dopamine, serotonin and norepinephrine neurons (Beckstead et al., 2009; Chen et al., 2012; Kirby et al., 2008; Kuhne et al., 2012; Lemos et al., 2012; Price et al., 2002; Riegel and Williams, 2008; Ungless et al., 2003; Valentino et al., 1983; Wanat et al., 2013; Wanat et al., 2008) as well as cholinergic neurons in basal forebrain, which mediates release of acetylcholine into the hippocampus (Day et al., 1998a). Prior stress history differentially affects the function of CRF at these different cellular substrates. For example, both stressor exposure as well as basal stress hyperresponsivity ablate the ability of CRF-R1 to increase GABAergic transmission onto serotonergic dorsal raphe neurons (Lamy and Beck, 2010; Lemos et al., 2011), thereby reducing CRF mediated-inhibition of serotonin release. Likewise, different types of stressor were shown to disrupt CRF’s ability to enhance dopamine transmission both in the VTA and the NAc (Beckstead et al., 2009; Lemos et al., 2012), an effect that our findings here confirm and extend upon. In contrast, stressor exposure potentiates the CRF-mediated increase in noradrenergic LC firing rates, particularly at lower concentrations, and potentiates the CRF-mediated increase in acetycholine release in the hippocampus (Day et al., 1998a).

Understanding the precise changes to the CRF system caused by stress exposure as well as the underlying mechanism by which this take place, has been complicated by the fact the direct comparison across different stress paradigms, species and preparations are not possible. In this study, we compared the effect of exogenously applied CRF at a given concentration (100 nM) on two different CRF effectors within the same animals in the same region. We found that while one effector is clearly disrupted following stressor exposure, the other is kept intact. These data support the notion that stress-induced plasticity of CRF function occurs in cell-type specific manner presumably to adaptively shift behavior responsivity. This main conclusion is consistent with recent work demonstrating that CRF-R1 deletion selectively in dopamine neurons leads to anxiogenic-like behaviors while CRF-R1 deletion in glutamatergic neurons produces anxiolytic-like responses (Refojo et al., 2011).

Collectively, the broad distribution of CRF-R1 expression to different cell-types, the different functional and behavioral consequences of CRF-R1 activation and the different sensitivity to stress-exposure by the individual CRF targets all point to the overall conclusion that the CRF system is quite complex and CRF release in different brain regions would trigger different behavioral responses, even when all mediated by the same receptor type, CRF-R1 in this case. In light of these data, it is logical to predict that no classical pharmacological manipulation that targets all CRF-R1 receptors would be ideal for treating stress-induced mood disorders. We propose that targets downstream of the CRF-R1 activation could offer more selectivity. For example, the M5 receptor is an interesting target for potential therapeutic actions, which unlike other muscarinic receptors it has quite restricted expression. Our data suggests that deletion of M5 receptors very selectively impairs the CRF actions on dopamine transmission. It is possible that boosting M5 receptor activity, e.g. with a positive allosteric modulator or partial agonist, would have therapeutic value by ameliorating some stress-induce impairments of dopamine-dependent behaviors such as exploratory and appetitive behaviors.

### Future Directions

There are many new avenues to pursue and follow up on from this first study. Perhaps the most pressing and obvious question is how CRF ultimately affect the excitability, firing and output of medium spiny projection neurons, which make up 95% of striatal neurons and constitute the output of sensory, motor and limbic integration from the striatum. There have been two reports that indicate that CRF enhances both c-fos immunoreactivity and phospho-CREB activity in striatal neurons (Pecina et al., 2006; Stern et al., 2011). It is still unknown (1) whether this connection is mediated through neuromodulation by DA and/or ACh, (2) whether this leads to changes in excitability and output of MSNs, and (3) finally whether it is selective for direct or indirect pathway MSNs. These are the next critical questions, which answers will further our understanding of CRF modulation of striatal circuit function and its behaviors.

## Author Contributions

Conceptualization: J.C.L. and V.A.A., Methodology: J.C.L, J.H.S., and V.A.A., Investigation, validation and analysis: J.C.L., J.H.S, A.I., and L.K.D. Writing manuscript:: J.C.L and V.A.A., Funding: V.A.A. and J.C.L., Resources: V.A.A. and J.C.L. Supervision: V.A.A. and J.C.L.

## Acknowledgements

This study was funded by the Intramural Programs of NIAAA, NINDS (ZIA-AA000421) to VAA, K99/R00 Pathway to Independence award (MH109627) to JCL and 2017 Innovation Award from NIH-DDIR to VAA. We thank Dr. Jurgen Wess for the M5KO mice. We are grateful to Dr. Steven Vogel for access to the confocal microscope. We thank Dr. David Lovinger and other members of the Alvarez lab for helpful comments and discussion. We also thank K. Deisseroth (Stanford) who generously provided the ChR2 construct.

## Experimental Procedures

All procedures were performed in accordance with guidelines from the Animal Care and Use Committee of the National Institute on Alcohol Abuse and Alcoholism.

### Animals

Male mice (p60-180) were group housed and kept under a 12h-light cycle (6:30 ON/18:30 OFF) with food and water available *ad libitum*. For *ex vivo* electrophysiology experiments and repeated stressor exposure followed by electrophysiology and voltammetry experiments, we used mice that were a cross between *ChAT-ires-Cre^+/−^ (Jax stock no.: 018957) × Ai14;tdTomato Reporter (B6.* Cg-Gt(ROSA)26Sortm14(CAG-tdTomato)Hze/J, Jax stock no.: 007914). M5KO homozygous mice backcrossed onto a C57BL6/J background were provided by Dr. Jurgen Wess and bred in the vivarium facility at NIAAA. Control mice were C57BL6/J mice obtained from Jackson Laboratories and bred concurrently in the vivarium facility at NIAAA. C57 control mice and M5KO mice were age matched.

### Fluorescent *In Situ* Hybridization (ISH) using RNAscope ®

Brains were rapidly dissected and flash frozen in isopentane on dry ice. Brain were kept in a −80°C freezer until they were sectioned. Coronal or sagittal brain slices (16 µm) containing the DS and NAc were thaw mounted onto Superfrost plus slides (Electron Microscopy Sciences) utilizing a Leica CM 1900 cryostat maintained at −20°C. Note, prior to section, brains were equilibrated in the cryostat for at least 2 hrs. Slides were cleaned with RNAzap, to prevent mRNA degradation. Slides were stored at −80°C.

RNAscope ISH was conducted according to the Advanced Cell Diagnostics user manual and as previously reported (Tejeda 2013, 2017). Briefly, slides were fixed in 10% neutral buffered formalin for 20 min at 4°C. Slides were washed 2×1 min with 1x PBS, before dehydration with 50% ethanol (1 × 5 min), 70% ethanol (1 x 5 min), and 100% ethanol (2 x 5 min). Slides were incubated in 100% ethanol at −20°C overnight. The following day, slides were dried at room temperature (RT) for 10 min. A hydrophobic barrier was drawn around the sections using a hydrophobic pen and allowed to dry for 15 min at RT. Sections were then incubated with Protease Pretreat-4 solution for 20 min at RT. Slides were washed with ddH2O (2 x 1 min), before being incubated with the appropriate probes for 2 hr at 40°C in the HybEZ oven (Advanced Cell Diagnostics). The following probes were purchased from Advanced Cell Diagnostics: Mm-*Crh1-* C1 (ACD Cat no: 418011), Mm-*Crh2-* C1 (ACD Cat no: 413201), Mm-*Crh*-C1 (ACD Cat no: 316091), and Mm-*Chat*-C2 (ACD Cat no:408731-C2) and Mm-*Th*-C3 (ACD Cat no: 317621-C3). Following incubation with the appropriate probes, slides were subjected to a series of amplification steps at 40°C in the HybEZ oven with 2 × 2 min washes (w/ agitation) in between each amplification step at RT. Amplification steps were as followed: Amp 1 at 40°C for 30 min. Amp 2 at 40°C for 15 min. Amp 3 at 40°C for 30 min. Amp 4-Alt A at 40°C for 15 min. A DAPI-containing solution was applied to sections (one slide at a time) at RT for 20 sec. Finally, slides were coverslipped using ProLong Gold Antifade mounting media and stored at 4°C until imaging on a confocal microscope (Zeiss).

### Image analysis and quantification

Sections were imaged using a Zeiss confocal microscope and Zen software. Unique 20x (5 µm thick) and 40x images (2 µm thick) were acquired from DS, NAc and VTA of stress-naïve and stress-exposed mice using the same software and hardware settings. The settings were titrated for each specific experimental probe. Quantification was done using Fiji/ImageJ software. Numbers of DAPI cells were automatically generated using the particle counter function in ImageJ. Numbers of ChAT cells and ChAT+/CRF-R1, +R2 or +CRF positive cells were manually counted using the cell counter function. For both ChAT and DAPI cells, cells were considered positive for the experimental probe, if there >5 particles clustered around (but not in) the cell nucleus. Numbers of puncta per ChAT (67-71 unique cells for naïve and stress-exposed mice) or TH (160 unique cells for naïve and stress-exposed mice each) cell was analyzed in a semi-automated fashion and this process is represented in SF9. Masks were generated using the ChAT or TH positive cell as a reference point. Rather than using the outline of the cell itself, we used a uniform circle that was the approximate size of the soma and this was kept consistent throughout the analysis. The fluorescent image of the experiment probe, in this case Mm-Crh1 or Mm-Crh, was converted into a binary image after being thresholded. The thresholding was kept consistent across images. Finally, the mask generated with the cell maker image was combined with the binary image of the experimental probe. This generated an image with only the puncta around the soma of our cells of interest were present. We then used the particle analyzer function to automatically count number of puncta.

### *In vitro* electrophysiology

Coronal slices (240 µm) containing the DS and NAc core were prepared from 8-16 week-old mice *ChAT-ires-CRE^+/−^;Ai14 tdTomato* mice. Slices were cut in ice-cold cutting solution (in mM): 225 sucrose, 13.9 NaCl, 26.2 NaHCO_3_, 1 NaH_2_PO_4_, 1.25 glucose, 2.5 KCl, 0.1 CaCl_2_, 4.9 MgCl_2_, and 3 kynurenic acid. Slices were maintained in oxygenated ACSF containing (in mM): 124 NaCl, 2.5 KCl, 2.5 CaCl_2_, 1.3 MgCl_2_, 26.2 NaHCO_3_, 1 NaH_2_PO_4_, and 20 glucose (∼310-315 mOsm) at RT following a 1hr recovery period at 33¼C. Cell-attached recordings in voltage clamp mode were performed to measure the effect of CRF on CIN firing frequency. For cell-attached recordings, electrodes were filled with filtered ACSF (the same that was used for external solution). A gigaohm seal was achieved, maintained and monitored. Cells in which the gigaohm seal had degraded were excluded. Current clamp recordings were performed using electrodes (2.5-4 MΩ) filled with 120 KMeSO_4_, 20 KCl, 10 HEPES, 0.2 K-EGTA, 2 MgCl_2_, 4 NaATP, and 0.4 Na-GTP (pH = 7.25, ∼290 mOsm). Data were acquired at 5 kHz and filtered at 1 kHz using Multiclamp 700B (Molecular Devices). Data was analyzed using pClamp (Clampfit, v. 10.3).

### Fast scanning cyclic voltammetry

Brain slices from C57BL6/J, M5KOor *ChAT-ires-CRE^+/^;Ai14 tdTomato* mice were prepared as described in the electrophysiology section using the same external solution and cutting solution. Carbon fiber (7 μm diam., Goodfellow) electrodes were fabricated with glass capillary (602000, A-M Systems) using a Sutter P-97 puller and fiber tips were hand cut to 100-150 μm past the capillary tip. Immediately prior to the experiments they were filled with 3M KCl internal solution. The carbon-fiber electrode was held at −0.4 V and a voltage ramp to and from 1.2 V (400V/s) was delivered every 100 ms (10 Hz). Before recording, electrodes were conditioned by running the ramp at 60 Hz for 15 min and at 10 Hz for another 15 min and calibrated. Electrodes were calibrated using 1 µM DA. DA transients were evoked by electrical stimulation delivered through a glass microelectrode filled with ACSF. Either a single monophasic pulse (0.2 msec, 300 µA) or trains of pulses (5 pulses at 5, 10, 25, 100 Hz) was delivered to the slice in the absence or presence of the nAChR antagonist DhβE (1 µM). For optogenetic experiments, light stimulation was delivered through optical fiber (0.6 msec, ∼700 µW). Data was acquired with a retrofitted headstage and Axon Amplifier using Multiclamp. Voltammetric analysis was done using custom written procedures in Igor Pro software. The voltamogram, peak amplitude, and area of the DA transient were measured. Experiments were rejected when the evoked current did not have the characteristic electrochemical signature of DA assessed by a current-voltage plot.

### Behavior

#### Forced Swim Stress

Mice were moved to behavioral suite and allowed to acclimate for at least 30 min. The swim stress procedure was done as described in (Lemos et al., 2012). Briefly, animals are placed in a 20 cm cylinder filled with 4L of water maintained at 30 ± 1¼C for 15 mins on Day 1 and 4 × 6 min on Day 2, separated by 6 min intervals in their homecage. Animals are returned to their homecage for 7 days and then *ex vivo* slices are prepared.

#### Statistics

Statistical analysis was performed in Prism (GraphPad) and Excel. Unless stated, two-tailed unpaired t-test was used. Otherwise, two-tailed paired t-test, 1W ANOVAs or 2W Repeated Measures (2WRM) ANOVAs or one-tailed t-tests were used when appropriate and stated. 2W ANOVAs were followed up with a Dunnett’s or Sidak-corrected t-test comparisons. All data are presented as mean ± SEM. Results were considered significant at an alpha of 0.05.

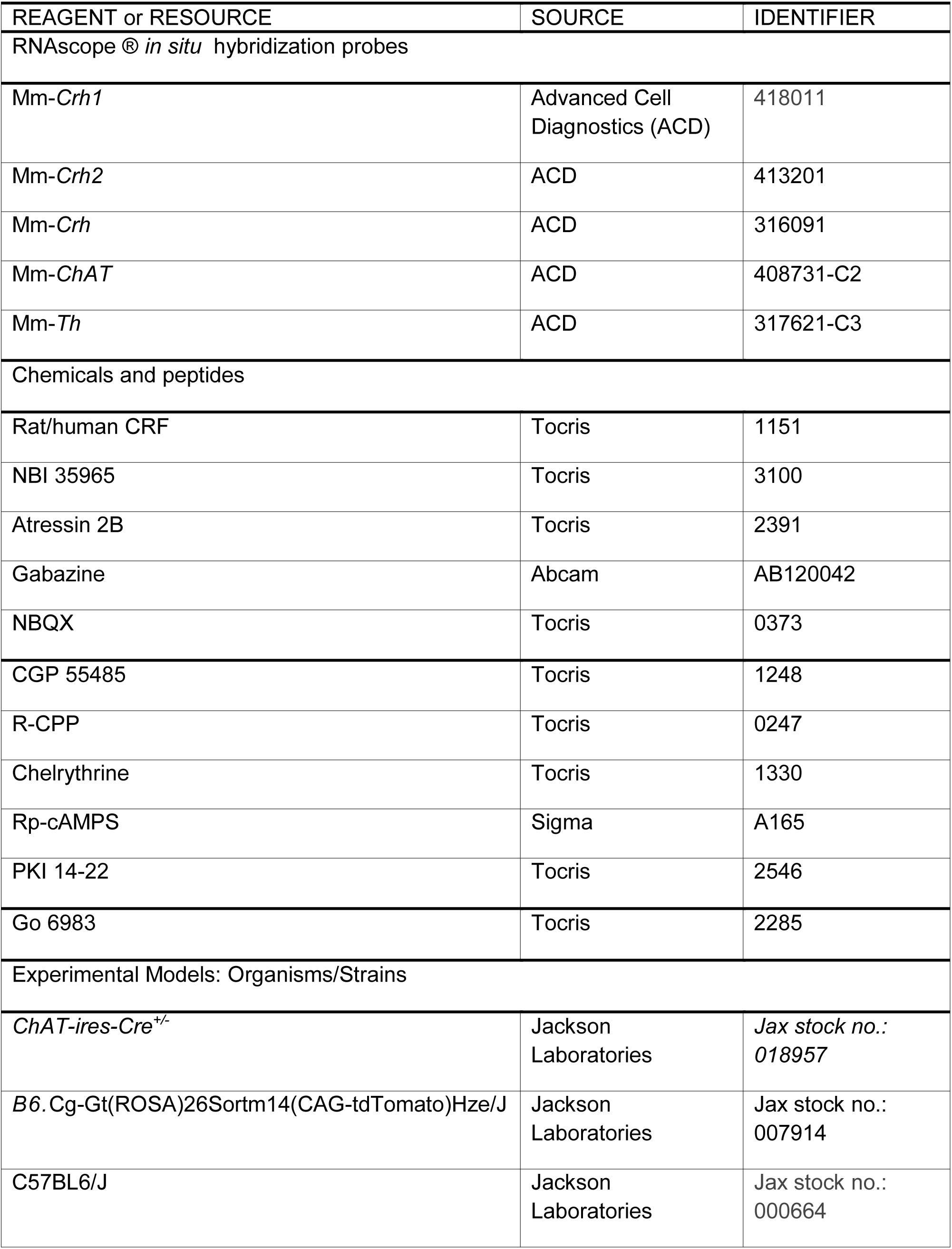
KEY RESOURCES TABLE.

